# A high throughput RNA displacement assay for screening SARS-CoV-2 nsp10-nsp16 complex towards developing therapeutics for COVID-19

**DOI:** 10.1101/2020.10.14.340034

**Authors:** Sumera Perveen, Aliakbar Khalili Yazdi, Kanchan Devkota, Fengling Li, Pegah Ghiabi, Taraneh Hajian, Peter Loppnau, Albina Bolotokova, Masoud Vedadi

## Abstract

SARS-CoV-2, the coronavirus that causes COVID-19, evades the human immune system by capping its RNA. This process protects the viral RNA and is essential for its replication. Multiple viral proteins are involved in this RNA capping process including the nonstructural protein 16 (nsp16) which is an S-adenosyl-L-methionine (SAM)-dependent 2’-O-methyltransferase. Nsp16 is significantly active when in complex with another nonstructural protein, nsp10, which plays a key role in its stability and activity. Here we report the development of a fluorescence polarization (FP)-based RNA displacement assay for nsp10-nsp16 complex in 384-well format with a Z′-Factor of 0.6, suitable for high throughput screening. In this process, we purified the nsp10-nsp16 complex to higher than 95% purity and confirmed its binding to the methyl donor SAM, product of the reaction, SAH, and a common methyltransferase inhibitor, sinefungin using Isothermal Titration Calorimetry (ITC). The assay was further validated by screening a library of 1124 drug-like compounds. This assay provides a cost-effective high throughput method for screening nsp10-nsp16 complex for RNA-competitive inhibitors towards developing COVID-19 therapeutics.

## Introduction

Over the last two decades, major outbreaks of coronaviruses have been endured worldwide. These include zoonotic epidemics of severe acute respiratory syndrome coronavirus (SARS-CoV) in 2002,^1^ Middle East Respiratory Syndrome coronavirus (MERS-CoV) in 2012,^2, 3^ and the current pandemic of SARS-CoV-2 that started in 2019 (COVID-19).^4^ With more than 34 million confirmed COVID-19 infections and a million deaths so far (October 3^rd^ 2020; World Health Organization; https://www.who.int/emergencies/diseases/novel-coronavirus-2019), the COVID-19 pandemic had widespread global socioeconomic implications, and overwhelmed health systems world-wide.

Coronavirus (CoV) is a member of the subfamily Coronavirinae within the family Coronaviridae.^5, 6^ Many species can be infected by CoVs.^6^ However, only seven coronaviruses have been documented to infect humans to date.^4, 7^ These include SARS-CoV, MERS-CoV, and SARS-CoV-2 which could cause severe symptoms leading to higher fatalities.^4^ Infection of 229E, HKU1, OC43, and NL63 has been associated with a range of relatively mild respiratory diseases.^7^ SARS-CoV-2 shares 79.6% RNA sequence identity with SARS-CoV.^8^ Coronaviruses are enveloped, non-segmented positive-sense RNA viruses which have the largest genome among RNA viruses ^9, 10^. Their genomes contain six major open reading frames (ORFs) and various number of accessory genes. They encode 16 nonstructural proteins (nsps) that play essential roles in RNA replication and processing of sub genomic RNA.^11–14^

SARS-CoV-2 replication involves RNA synthesis, proofreading and capping.^15^ Coronavirus mRNAs are protected at their 5′ ends by a cap structure consisting of an N7-methylated guanine linked to the first transcribed nucleotide by a 5′–5′ triphosphate bond and 2′-O-methylation (N7-meGpppN-2′-O-me; Fig. 1).^16^ The RNA capping is essential for stability of viral mRNA and evading the host immune system. Uncapped RNA molecules are identified as ‘foreign’ by host innate immune response and undergo degradation.^17, 18^ Therefore, proteins catalyzing RNA-capping are attractive targets for antiviral drug development. Nsps involved in viral mRNA capping include nsp10 (cofactor for nsp14 and nsp16),^19^ nsp13 (RNA 5’-triphosphatase, helicase),^20^ nsp14 (guanine-N7 methyltransferase),^21^ and nsp16 (2’-O-methyltransferase).^18, 22, 23^

**Figure 1.**
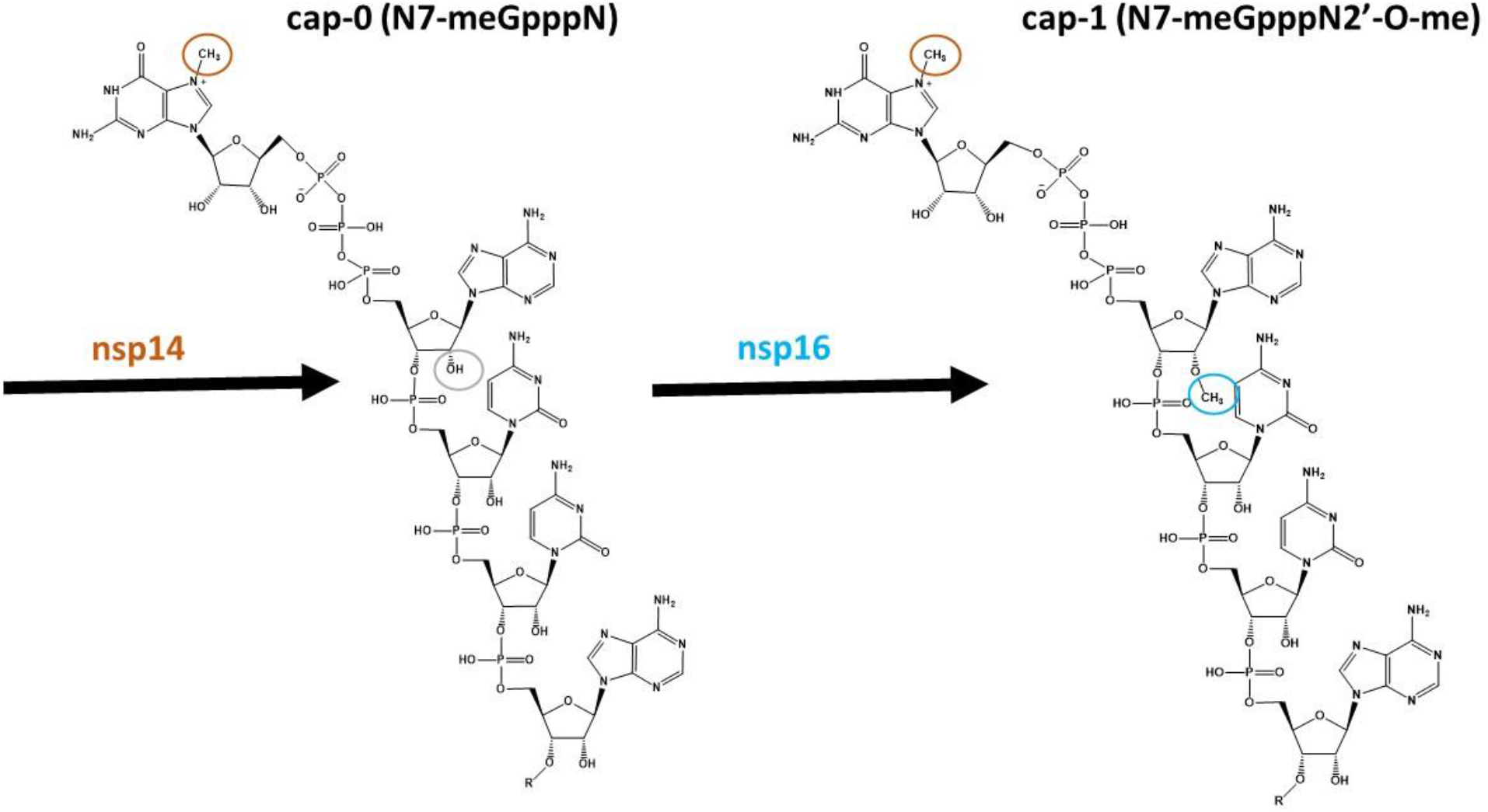
RNA capping by nsp14 and nsp16. Nsp14 catalyzes the formation of cap-0 (N7-meGpppN-RNA) by methylating N7 of the guanosine (brown circle). In turn, nsp16 converts cap-0 to cap-1 (N7-meGpppNme-RNA) by 2′-O-methylating (blue circle) the cap. Modified from the enzyme reactions presented in Brenda (www.brenda-enzymes.org).

The capping process begins with the removal of the 5’γ-phosphate of the newly synthesized RNA chains (pppN-RNA) by nsp13.^20^ Then a guanylyltransferase (GTase) leads to the formation of GpppN-RNA, by transferring a Guanosine monophosphate (GMP) molecule to the 5’-diphosphate of the RNA chains (ppN-RNA). Nsp14 methylates the cap structure at the N7 position of the guanosine, resulting in the formation of cap-0 (N7-meGpppN-RNA).^21^ Ultimately, the nsp10-nsp16 complex catalyzes the addition of a methyl group on the ribose 2′-O position of the first transcribed nucleotide of the cap-0 (N7-meGpppN-RNA) to form a cap-1 (N7-meGpppNme-RNA) (Fig. 1).^23, 24^ *In vitro*, methyltransferase activity has been reported for nsp10-nsp16 complexes from feline CoV (FCoV),^25^ SARS-CoV,^23^ and MERS-CoV,^26^ and the binding of various RNAs to MERS-CoV have been assessed and used to rank-order substrates.^26^

Nsp10 acts as an allosteric activator of nsp16 and is essential for its stability and activity,^18, 27^ increasing its binding affinity for RNA substrate and SAM in both SARS and MERS-CoV.^18^ Various structures of nsp10-nsp16 from SARS-COV-2 have been recently deposited in Protein Data Bank (PDB) such as PDB IDs: 6W4H, 6W75, 6WJT, 6WKQ, 6WVN, 6WQ3 and 6WRZ.^28^ These include the SARS-CoV-2 nsp10-nsp16 in complex with SAM (6W75), SAH (6WJT), sinefungin (6WKQ) and in complex with both RNA (N7-meGpppA) and SAM (6WVN) or RNA (N7-meGpppA) and SAH (6WQ3 and 6WRZ).^28^ Along with previously available structures from SARS-CoV nsp10-nsp16 (PDB IDs: 2XYR, 2XYV, 2XYQ, 3R24)^18, 22^ and MERS-CoV nsp16 PDB ID: 5YN5), these structures provide vast amount of information on interaction of substrates with nsp16 and could be used in structure guided drug discovery (Fig. 2).

**Figure 2.**
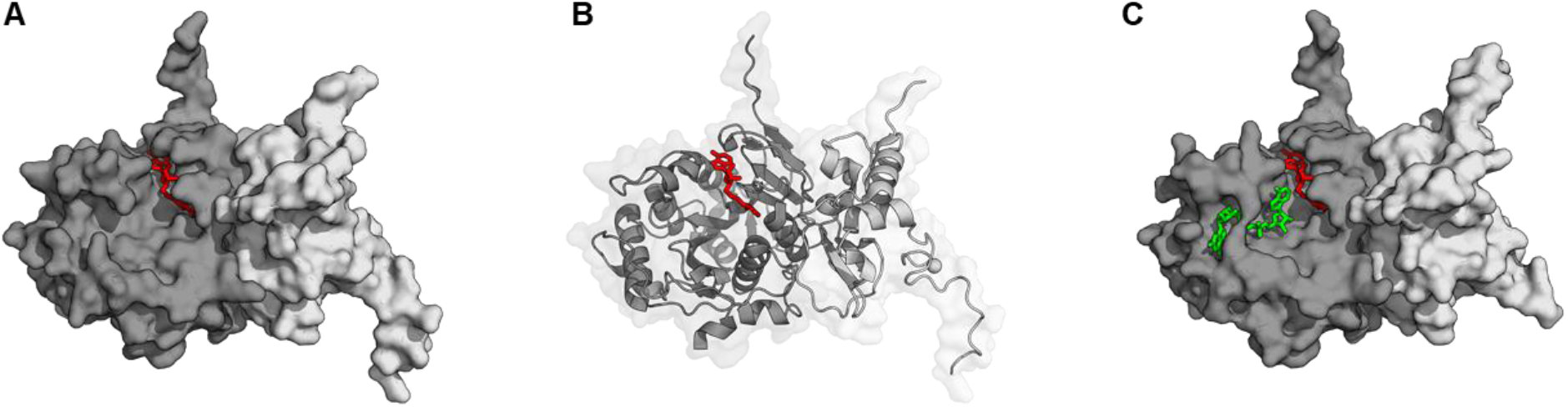
Crystal structures of SARS-CoV-2 nsp10-nsp16.^28^. Crystal structures of nsp10-nsp16 in complex with (A and B) SAH (PDB: 6WJT) and (C) N7-meGpppA and SAH (PDB: 6WQ3) have been recently reported and could be used in structure guided drug discovery. Nsp16 and nsp10 are in dark gray and light gray, respectively. SAH is shown with red sticks, while N7-meGpppA is represented in green.

Here we report the development of a cost-effective fluorescence polarization-based RNA-displacement assay which is suitable for high-throughput screening of nsp10-nsp16 complex against large libraries of compounds to identify RNA competitive inhibitors of nsp16 methyltransferase activity. This assay is also a perfect tool for triaging a high number of potential inhibitors from screening by alternative methods for competing with RNA substrate and determining their mechanism of inhibition.

## Materials and methods

### Reagents

S-adenosylmethionine (SAM), S-adenosylhomocysteine (SAH) and sinefungin were purchased from Sigma-Aldrich. FAM labelled RNA (5’ N7-meGpppACCCCC-FAM 3′) and biotinylated RNA (5’ N7-MeGpppACCCCC-Biotin 3′) were purchased from bioSYNTHESIS (Levisville, Texas). The unlabelled Cap RNA, m7G(5′)ppp(5′)G (NEB, #S1404S) and m7G(5′)ppp(5′)A (NEB, #S1405S) also referred throughout the manuscript as N7-meGpppG and N7-meGpppA for consistency, respectively, and G(5′)ppp(5′)A RNA (GpppA) (NEB, # S1406S) and G(5′)ppp(5′)G RNA (GpppG) (NEB, # S1407S) are from New England Biolabs (NEB). NEB RNA reagents were gifts from Dr. Peter Stogios, University of Toronto. All RNA solutions were prepared by dissolving in nuclease free water, in the presence of RNAseOUT (Life Technologies) at a final concentration of 0.4 U/μL.

### Protein Purification

Expression and purification of nsp10 and nsp16, and nsp10-nsp16 complex preparation have been described in detail in Supplementary data (Supplementary Fig. 1, 2 and 3).

### Fluorescence Polarization-based RNA displacement Assays

All fluorescence polarization (FP) experiments were performed in a total assay volume of 20 μL per well in 384-well black polypropylene PCR plates (Axygen, Tewksbury, MA, USA; cat. PCR-384-BK) using FAM (Fluorescein amidites) labelled RNA (5’ N7-meGpppACCCCC-FAM 3′) and nsp10-nsp16 complex. In all experiments, nsp10-nsp16 complex was used at a ratio of 8 (nsp10):1 (nsp16). Fluorescence polarization (FP) was measured after 30 min of incubation at room temperature, using a BioTek Synergy 4 (BioTek, Winooski, VT) with the excitation and emission wavelengths of 485 nm and 528 nm, respectively. All experiments were performed in triplicate (n=3) and plotted values are the average of 3 replicates ± standard deviation. The FP values were blank subtracted and were presented as the percentage of control (FP %). Data were visualized using GraphPad Prism software 7.04 (La Jolla, CA).

For K_d_ determination, varying concentrations of nsp10-nsp16 were incubated with 30 nM FAM-RNA (5’ N7-meGpppACCCCC-FAM 3′), in 10 mM Tris buffer pH 7.5 containing 5 mM DTT, 0.01% Triton X-100, 0.01% BSA for 30 min at room temperature in 20 μL reaction volume. Fluorescence polarization (485 nm/ 528 nm) was measured using a Biotek Synergy H1. Change in fluorescence polarization (mP) was plotted as a function of nsp10-nsp16 concentration. The binding K_d_ and maximal FP signal (Bmax) were calculated using nonlinear least squares regression to a single-site binding model in GraphPad Prism 7.04.

To establish the specificity of the assay, a mixture containing 0.5 μM nsp10-nsp16 complex and 30 nM FAM-RNA, in 10 mM Tris buffer pH 7.5 containing 5 mM DTT, 0.01% Triton X-100, 0.01% BSA were incubated separately with varying concentrations of unlabelled RNA cap analogues; N7-meGpppG, N7-meGpppA, GpppG and GpppA in 20 μL reaction volume respectively for 30 min at room temperature. The FP values were determined, and the K_disp_ (K displacement) values (the concentrations required for 50% displacement of the labeled RNA) were calculated using nonlinear least squares regression analysis to a four-parameter concentration-response curve model in GraphPad Prism 7.04.

### Z′ factor determination

The *Z*′-factor was determined by incubating nsp10-nsp16 complex (0.5 μM) with 30 nM FAM-RNA in the presence or absence of 50 μM unlabeled RNA (N7-meGpppA) in a 384-well plate for 30 min at room temperature (168 wells each). Fluorescence polarization was measured as described above.

### Screening compounds from the Prestwick library

Prestwick Chemical Library was purchased from Prestwick Chemicals and was screened using optimized assay conditions (10 mM Tris buffer pH 7.5 containing 5 mM DTT, 0.01% Triton X-100, 0.01% BSA) at 50 μM compound concentration, with 0.5% DMSO concentration in 384-well black polypropylene PCR plates (Axygen, Tewksbury, MA, USA; cat. PCR-384-BK) with a final reaction volume of 20 μL for 30 min at room temperature. Data were analyzed using GraphPad Prism 7.04 as described above.

### Isothermal Titration Calorimetry (ITC)

Binding of SAM, SAH, and sinefungin to nsp10-nsp16 (8:1) complex was tested by Nano ITC titration calorimeter (TA Instruments — Waters LLC, New Castle, DE) at 25 °C. Nsp10-nsp16 complex was dialyzed against 50 mM Tris-HCl buffer (pH 8.0) containing 200 mM NaCl and was loaded into the sample cell (300 μL) at a concentration of 48 μM, and a solution of 500 μM of ligand (SAM, SAH, or sinefungin) was placed in the injection syringe (50 μL). Data were fitted with a one binding site model using TA Instruments origin software.

## Results

### SARS-COV-2 nsp10-nsp16 complex preparation and quality control

His-tagged nsp10 (residues A1-Q139) and nsp16 (S1 – N298) were expressed in *E. coli* separately and purified to homogeneity using Ni-NTA resin as described in the supplementary material and methods (Supplementary Fig.1). Purified nsp10 and nsp16 proteins were mixed at a molar ratio of 8 to 1, respectively, to prepare stable and functional complex. The complex formation was evaluated by running a sample of expected complex through size exclusion chromatography which confirmed complex formation (Supplementary Fig. 2). In addition, the presence of intact nsp10 and nsp16 in the complex fractions were verified by mass spectrometry (Supplementary Fig. 3).

To evaluate the functional integrity of the prepared complex, we tested the binding of the methyl donor SAM, product of the reaction SAH, and a common methyltransferase inhibitor sinefungin using ITC. Nsp10-nsp16 complex showed binding to SAM, SAH and sinefungin with K_d_ values of 3.4 ± 1.5 μΜ, 5.7 ± 1.9 μΜ, and 6.8 ± 2.0 μΜ, respectively, indicating the protein complex was formed and maintained the expected active site conformation (Fig. 3).

**Figure 3.**
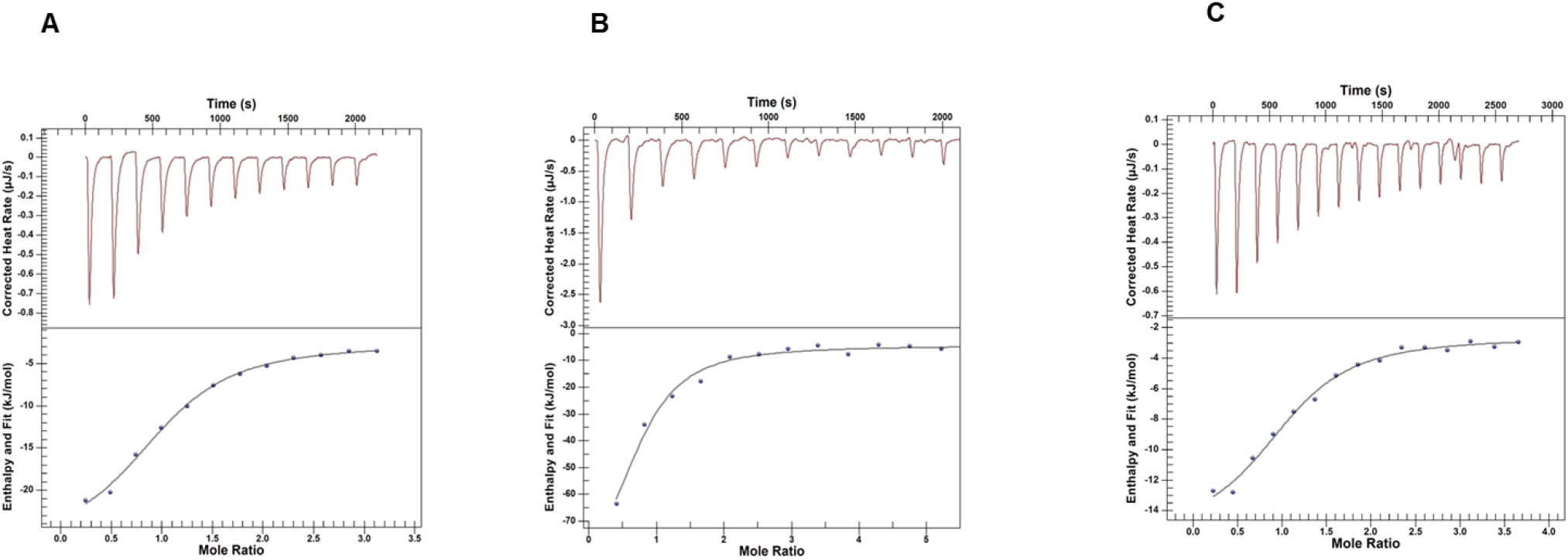
Isothermal Titration Calorimetry (ITC). ITC analysis of nsp10-nsp16 binding to (A) SAM, (B) SAH and (C) sinefungin with K_d_ values of 3.4 ± 1.5 μΜ, 5.7 ± 1.9 μΜ, and 6.8 ± 2.0 μΜ, respectively. The K_d_ values are mean ± s.d. of three independent experiments.

### RNA-displacement assay development

We used a FAM-labelled RNA (5’ N7-meGpppACCCCC-FAM 3′) to set up an RNA displacement assay. We first tested binding of this RNA to the nsp10-nsp16 complex. We observed significant fluorescence polarization (FP) signal and proceeded to optimize the assay conditions (Supplementary Fig. 4). Various buffer conditions such as 4-(2-hydroxyethyl)-1-piperazineethanesulfonic acid (HEPES), Bis-Tris Propane (BTP), Sodium phosphate (NaP), Tris(hydroxymethyl)aminomethane (Tris) and Potassium phosphate (KP) at 20 mM, pH 7.5 were tested and the FP signal was relatively higher in Tris and HEPES than other buffers (Supplementary Fig. 4A).

We selected to use 10 mM Tris for testing additives, which provided the highest signal (Supplementary Fig. 4B). The signal was unaffected up to 1% BSA, 10 mM of TCEP, KCl and MgCl_2_ (Supplementary Fig. 4C to 4F). The pH profile showed pH 7.5 as optimum (Fig. 4A). The signal was not significantly affected by increasing concentrations of DTT (Fig. 4B), and NaCl (Fig. 4C), or EDTA (Fig. 4D) up to 10 mM. Triton X-100 (Fig. 4E) and DMSO (Fig. 4F) had no significant effect on signal up to 1% and 5%, respectively. DTT was selected to be used at 5 mM in assays to maintain a reducing environment. Addition of Triton X-100 at 0.01% is expected to help minimizing binding of fluorophores and proteins to polystyrene plates ^29^ and compound aggregate formation in screening ^30^. BSA was also included at 0.01% to prevent non-specific interactions. Based on optimization results, the final assay buffer and screening conditions established consisted of 5 mM DTT, 0.01% Triton X-100, 0.01% BSA in 10 mM Tris buffer pH 7.5, and incubation time of 30 min.

**Figure 4.**
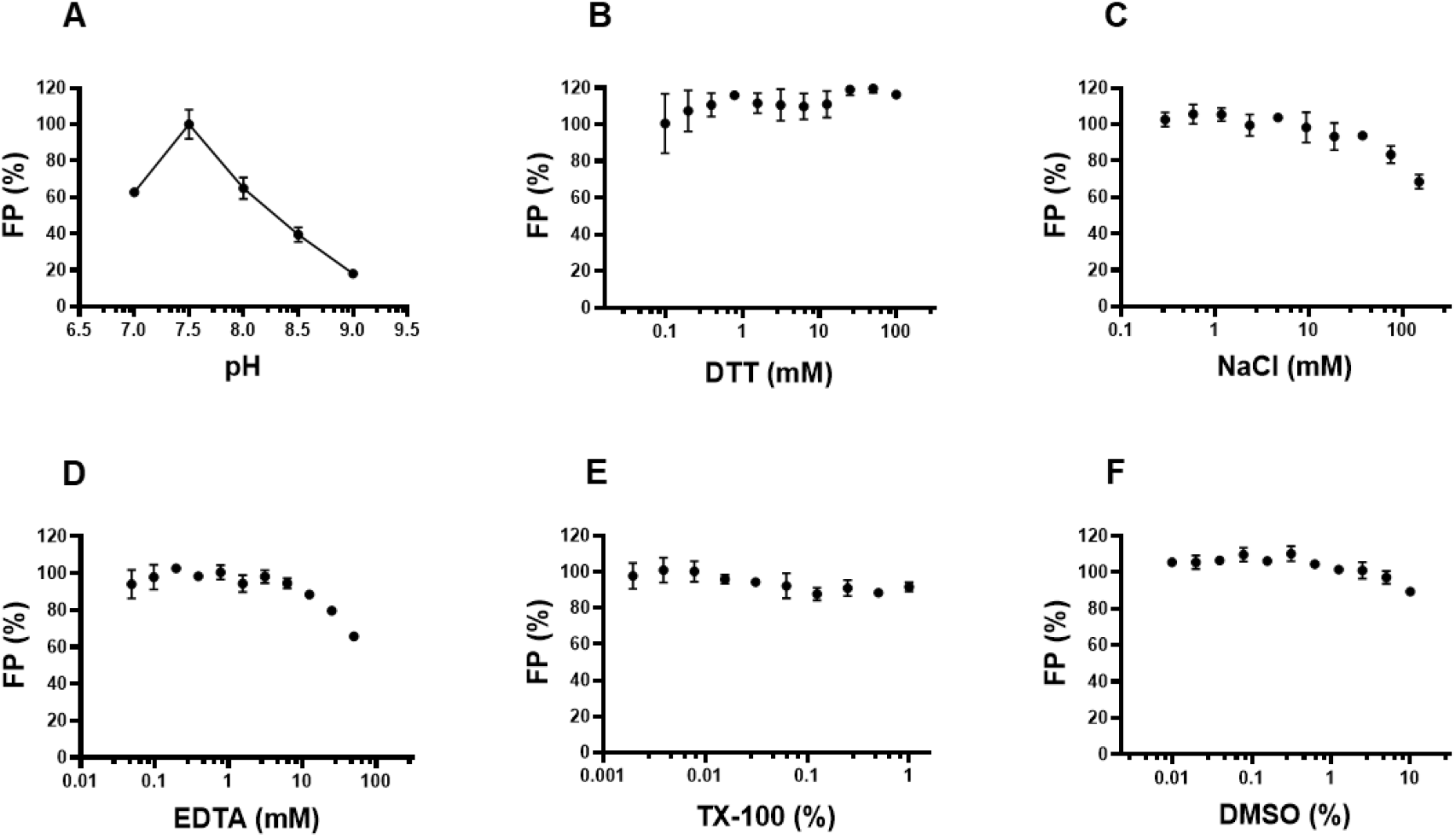
Fluorescence polarization (FP) assay optimization for nsp10-nsp16 interaction with N7-meGpppACCCCC-FAM. Fluorescence polarization signal from the interaction of 0.5 μM nsp10-nsp16 with 30 nM N7-meGpppACCCCC-FAM was evaluated (A) at pH range of 7.0 to 9.0 in 10 mM Tris buffer. Effect of additives such as (B) DTT; (C) NaCl (D) EDTA; (E) Triton X-100 and (F) DMSO was evaluated in 10 mM Tris, pH 7.5. DMSO had little effect on assay signal up to 5%. Experiments were performed in triplicate (n=3) and plotted values are the average of 3 replicates ± standard deviation. The FP values are blank subtracted and are presented as the percentage of control, in (A) pH 7.5 was considered as 100% signal. Data were analyzed using GraphPad Prism software 7.0.4.

Using the optimized assay conditions, nsp10-nsp16 complex showed concentration dependent binding to the FAM-RNA at 30 nM with an apparent K_d_ of 0.13 ± 0.002 μM and Bmax of 270 ± 5 mP (Fig. 5A). For RNA-displacement assays, the protein complex concentration was kept at 80% of maximum signal (0.5 μM) to allow better competitive displacement. Unlabelled methylated RNA cap analogues, N7-meGpppG and N7-meGpppA, were used to compete off the FAM-RNA binding to nsp10-nsp16 complex. Both N7-meGpppG and N7-meGpppA displaced FAM-RNA with K_disp_ values of 17 ± 3 μM and 20 ± 6 μM, respectively (Fig. 5B and 5C), while unlabelled unmethylated cap analogues GpppG (Fig. 5D) and GpppA (Fig. 5E) were unable to displace FAM-RNA from nsp10-nsp16 complex. C-terminally biotinylated 5′ N7-methylated GpppACCCCC 3′ also displaced the FAM-RNA with K_disp_ of 4.3 μM (Supplementary Fig. 5). These observations further confirmed that the nsp10-nsp16 protein complex specifically recognizes its N7-me RNA substrate.

**Figure 5.**
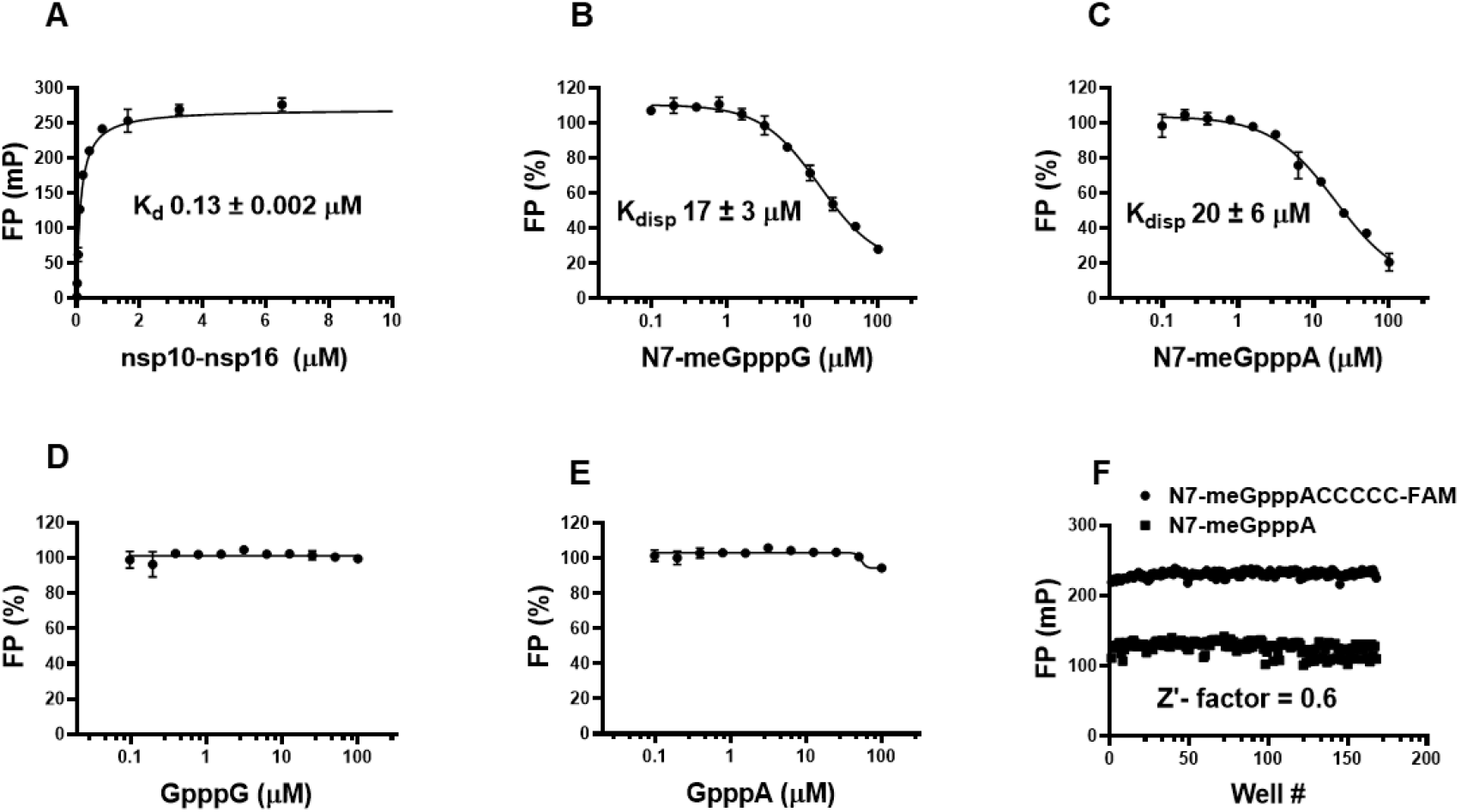
Fluorescence polarization (FP) saturation binding and competitive displacement curves for nsp10-nsp16 complex. (A) nsp10-nsp16 complex binds to 5’ N7-meGpppACCCCC-FAM 3′ with K_d_ of 0.13 ± 0.002 μM and Bmax of 270 ± 5 mP; Unlabeled methylated cap RNAs, (B) N7-meGpppG and (C) N7-meGpppA, compete with 5’ N7-meGpppACCCCC-FAM 3′ for binding to nsp10-nsp16 complex. Unlabeled unmethylated cap RNAs, (D) GpppG and (E) GpppA, did not disrupt the interaction between nsp10-nsp16 complex and RNA-FAM. All values are mean ± s.d. of three independent experiments. (F) Z’-factor (0.6) was determined for evaluation of the assay for high-throughput screening. FAM labelled RNA (5’ N7-meGpppACCCCC-FAM 3′) at 30 nM was used to generate the FP signal, while 50 μM unlabeled RNA (N7-meGpppA) was used as a positive control. Data was analyzed using GraphPad Prism software 7.0.4.

To determine whether the RNA-displacement assay at optimized conditions is amenable to high-throughput screening, the Z′-factor was determined by performing the assay in a 384-well format using FAM-RNA (5’ N7-meGpppACCCCC-FAM 3′) in the presence (168 data points) and absence (168 data points) of 50 μM unlabelled RNA (N7-meGpppA). The calculated Z′-factor was 0.6, indicating the assay is suitable for screening for RNA competitive inhibitors (Fig. 5F).

### Screening a library of 1124 compounds

We further screened a collection of 1124 small molecules from the Prestwick Chemical Library, a collection of drugs and drug-like compounds. Screening of this library initially identified 15 compounds as potential antagonists of nsp10-nsp16-RNA interaction based on reduction of polarization signal (>50% reduction) compared to the control at 50 μM final compound concentration (Fig. 6).

**Figure 6.**
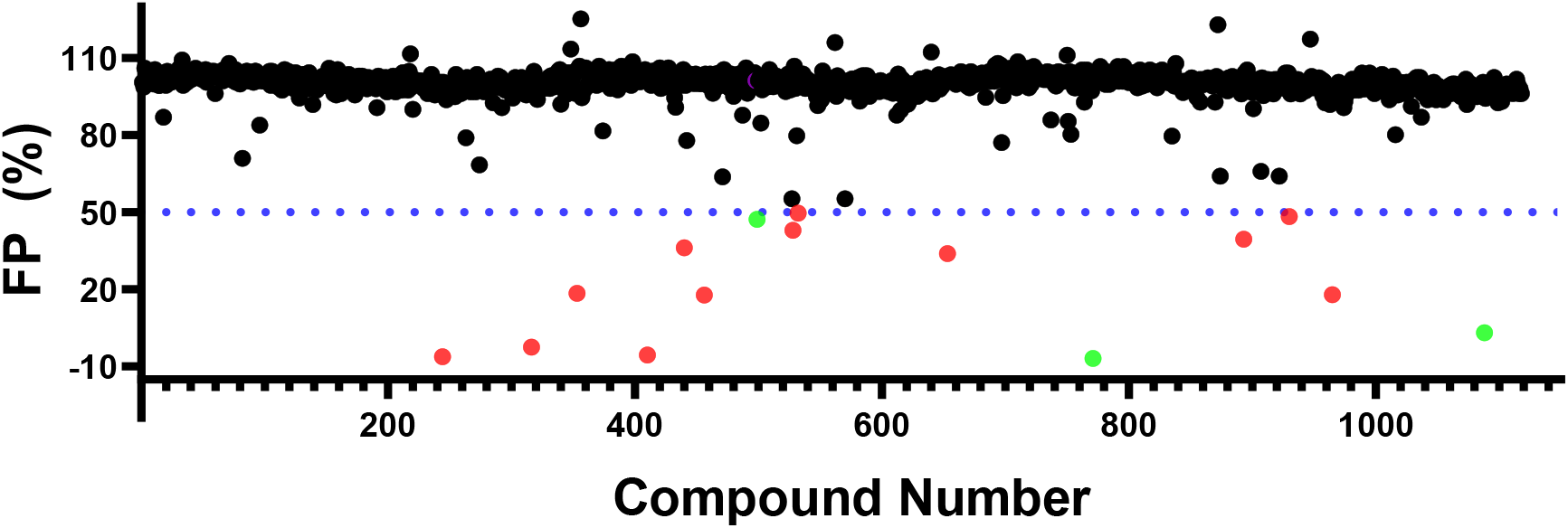
Screening of Prestwick compound Library: (A) Distribution of readout for 1124 compounds tested for binding to nsp10-nsp16 by FP-based RNA displacement assay. Compounds that showed less than 50% polarization signal compared to control were selected as screening hits (highlighted in red) for follow ups. Three compounds (highlighted in green) showed concentration dependent inhibition. However, none of these compounds showed any binding when tested by ITC.

All 15 compounds were tested for any possible interference with the signal readout, and 12 compounds showed a significant effect and were eliminated. Three compounds showed dose-dependent displacement with no signal interference. However, considering the high Hill slopes as a red flag, we further tested the binding of these 3 compounds to nsp10-nsp16 complex by ITC and none of the compounds was confirmed. The screening test run once again confirmed the suitability of the optimized RNA displacement assay for medium to high throughput screening of nsp10-nsp16 complex.

## Discussion

The COVID-19 pandemic has an unprecedented impact on the global economy and has already brought the health care services worldwide to a breaking point. So far, there are no approved vaccine or effective drugs for this disease. Even vaccines developed today may not be effective on future mutated variants.^31^ This necessitates the development of therapeutics that could stop the replication of the SARS-CoV-2. Coronavirus caps its RNA to evade human immune system and preserve its RNA from degradation, which is essential for viral replication.^15, 32^ RNA-capping is catalyzed by non-structural enzymes including nsp16 which is a SAM-dependent RNA-methyltransferase, and active when in complex with nsp10.^26^ Methyltransferases have been shown to be druggable^33^ with more than 20 potent, selective, and cell active small molecules (chemical probes) discovered for such enzymes in the last decade^33^. Typically, both SAM^34^ and substrate binding sites of methyltransferases could be targeted for drug discovery.^33^ One of the cost-effective ways of screening for RNA substrate competitive inhibitors is to label the RNA with fluorescein and detect its binding to the target RNA methyltransferase by monitoring the change in fluorescent polarization signal.^35^ In this study, we have developed and optimized a fluorescence polarization (FP)-based RNA displacement assay in a 384-well format for nsp10-nsp16 complex which is suitable for screening to identify RNA-competitive inhibitors. This assay is also suitable for triaging RNA-competitive small molecules from a high number of hits that could be identified by other types of screening.

SAM has been reported to promote active MERS nsp10-nsp16 complex assembly and SAM/SAH balance regulating nsp16 activity.^26^ The K_d_ values reported for SAM binding to SARS-CoV nsp10-nsp16 complex (K_d_ = 5.59 ± 1.15 μΜ) ^22^ are within the same range as we report for SARS-CoV-2 nsp10-nsp16 complex binding to SAM (K_d_ = 3.4 ± 1.5 μM). This indicates the highly conserved interaction of co-factor SAM with nsp10-nsp16 from these coronaviruses. The FAM labelled RNA substrate in this study (5’ N7-meGpppACCCCC-FAM 3′) was designed based on the previous studies on SARS-CoV nsp10-nsp16 complex reporting recognition of cap-0 RNA (N7-meGpppA-RNA) as substrate but not N7-meGpppG-capped RNA.^22^ In vitro, RNA substrate for SARS-CoV nsp10-nsp16 needs to be N7 guanine methylated with adenine (A) as the first nucleotide ^23^. We observed no RNA displacement with SARS-CoV-2 nsp10-nsp16 using the unmethylated RNA samples such as GpppA and GpppG. These observations are consistent with previous reports on similar binding assays with SARS-CoV^18^ and MERS-CoV^26^ proteins, where nsp16 can discriminate the cap-0 (N7-meGpppN-RNA) structure from an unmethylated cap structure (N7-GpppN-RNA). Nsp10-nsp16 complex was reported active as a type-0 RNA cap-dependent 2’-O-methyltransferasease.^23^

The RNA-displacement assay in a 384-well format presented in this study provides a cost-effective method to screen nsp10-nsp16 complex against large libraries of compounds to identify starting small molecule RNA-competitive inhibitors. Such molecules could lead to developing therapeutics for COVID-19 through inhibition of viral replication, and likely would be effective on other coronaviruses due to high conservation of nsp16 among this family of viruses.

## Acknowledgements

We thank Dr. Matthieu Schapira for helpful discussions, Dr. Peter Stogios for providing reagents, Dr. Peter Brown for review of the manuscript, and Dr. Aled Edwards and Dr. Cheryl Arrowsmith for continued support. This research was funded by the University of Toronto COVID-19 Action Initiative-2020, Takeda California, Inc., and COVID-19 Mitacs Accelerate postdoctoral awards to AKY and SP. The Structural Genomics Consortium is a registered charity (no: 1097737) that receives funds from; AbbVie, Bayer Pharma AG, Boehringer Ingelheim, Canada Foundation for Innovation, Eshelman Institute for Innovation, Genentech, Genome Canada through Ontario Genomics Institute [OGI-196], EU/EFPIA/OICR/McGill/KTH, Diamond Innovative Medicines Initiative 2 Joint Undertaking [EUbOPEN grant 875510], Janssen, Merck KGaA (aka EMD in Canada and US), Merck & Co (aka MSD outside Canada and US), Pfizer, São Paulo Research Foundation-FAPESP, Takeda and Wellcome [106169/ZZ14/Z].

## Author contributions

S.P designed and performed experiments, analyzed data and wrote the manuscript, A.K.Y contributed to the manuscript preparation, K.D performed Prestwick library screening, F.L performed ITC experiments, P.G and T.H purified proteins and prepared the protein complex, P.L. cloned constructs, A.B contributed to compound management, M.V was the Principal Investigator and lead the overall project, designed experiments, reviewed data and wrote the manuscript with input from all authors.

## Protein expression and purification

DNA sequences encoding the full-length SARS-CoV-2 nsp10 (residues A4254-Q4392 / A1-Q139) and nsp16 (residues S6799-N7096 / S1-N298) were chemically synthesized according to the UniProtKB reference sequence P0DTD1. The DNA fragments were individually cloned into the pET28-MHL expression vector (GenBank: EF456735.1) adding a N-terminal purification tag (MHHHHHHSSGRENLYFQG).

Constructs were transformed into *Escherichia coli* BL21 (DE3) RIL, then the cells were cultured in LB Broth overnight at 37 °C. The next day, the overnight cultures were used to inoculate Terrific Broth supplemented with 50 μg/mL Kanamycin and 35 μg/mL chloramphenicol for further amplification at 37 °C using LEX system (https://www.thesgc.org/science/lex). Once the OD600 of the cultures reached 0.8-1.5, the temperature was lowered to 18 °C and the target proteins were overexpressed overnight by inducing cells with 1 mM IPTG (isopropyl-1-thio-D-galactopyranoside). The cells were harvested the next day at 7000 rpm for 10 minutes at 4 °C (Beckman Coulter centrifuge).

Harvested cells from nsp10 and nsp16 cultures were re-suspended in binding buffer containing 50 mM Tris-HCl pH 8.0, 500 mM NaCl, 5% Glycerol, 0.5 mM TCEP, 5 mM imidazole and 50 mM Tris-HCl pH 9.0, 500 mM NaCl, 5% Glycerol, 0.5 mM TCEP, 5 mM imidazole, respectively supplemented with 1X protease inhibitor cocktail (100X protease inhibitor stock in 70% ethanol containing 0.25 mg/mL Aprotinin, 0.25 mg/mL Leupeptin, 0.25 mg/mL Pepstatin A and 0.25 mg/mL E-64) or Roche complete EDTA-free protease inhibitor cocktail tablet. The cells were lysed chemically by rotating 30 min with CHAPS (final concentration of 0.5%) and 5 μL/L Benzonase Nuclease (in house) followed by sonication at frequency of 8.0 (10s on/7s off) for 5 min (Sonicator 3000, Misoni). The crude extract was clarified by high-speed centrifugation for 45 min at 16000 rpm at 4 °C (Beckman Coulter centrifuge).

The lysates were then separately loaded onto Ni-NTA affinity resin column (Qiagen) pre-equilibrated with the corresponding binding buffers. The column for nsp10 was washed with running wash buffer (binding buffer plus 30 mM imidazole). The column for nsp16 was once washed by wash buffer and 4 more times with binding buffer. Subsequently, the columns were eluted with the corresponding binding buffers containing 250 mM imidazole (Fig. S1). Then, the nsp16 and nsp10 proteins were mixed at 1:8 molar ratio and the mixture was dialyzed in final buffer containing 50 mM Tris-HCl pH 8.0, 200 mM NaCl, 5% glycerol, 0.5 mM TCEP for a total of 60 minutes while spinning at 4 °C. Then, the complex was flash frozen and stored at −80 °C.

We evaluated nsp10-nsp16 complex formation by Size-Exclusion Chromatography (SEC). Initially, a SEC standard (Bio-Rad) was loaded onto Superdex 200 16/600 column after equilibration with 50 mM Tris-HCl pH 8.0 buffer containing 200 mM NaCl, 5% glycerol, 0.5 mM TCEP. The standard peaks were used to plot the calibration curve. Then, the protein sample was loaded onto the column and the molecular weight of the protein fractions were calculated according to the calibration curve (Fig. S2). The molecular weight of the nsp10 and nsp16 was confirmed by running 10 μg of protein complex on mass spectrometer (Agilent Technologies, 6545 Q-TOF LC/MS) (Fig. S3).

---

**Figure S1.**
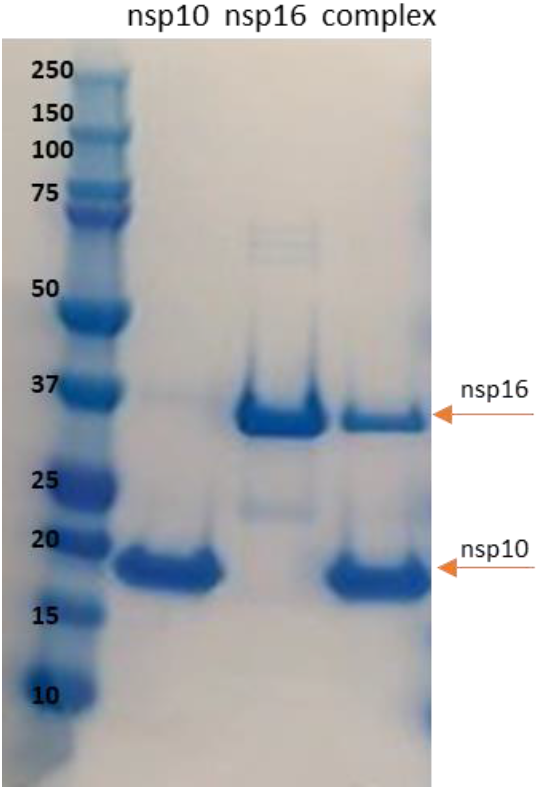
Purification of nsp10 and nsp16. SDS-PAGE electrophoresis (4-12% Bis-Tris gel; Life Technologies) demonstrating Ni-NTA purification of nsp10 and nsp16, and the nsp10-nsp16 complex preparation. The arrows show nsp16 and nsp10 at approximately 35.5 and 16.9 kDa, respectively.

**Figure S2.**
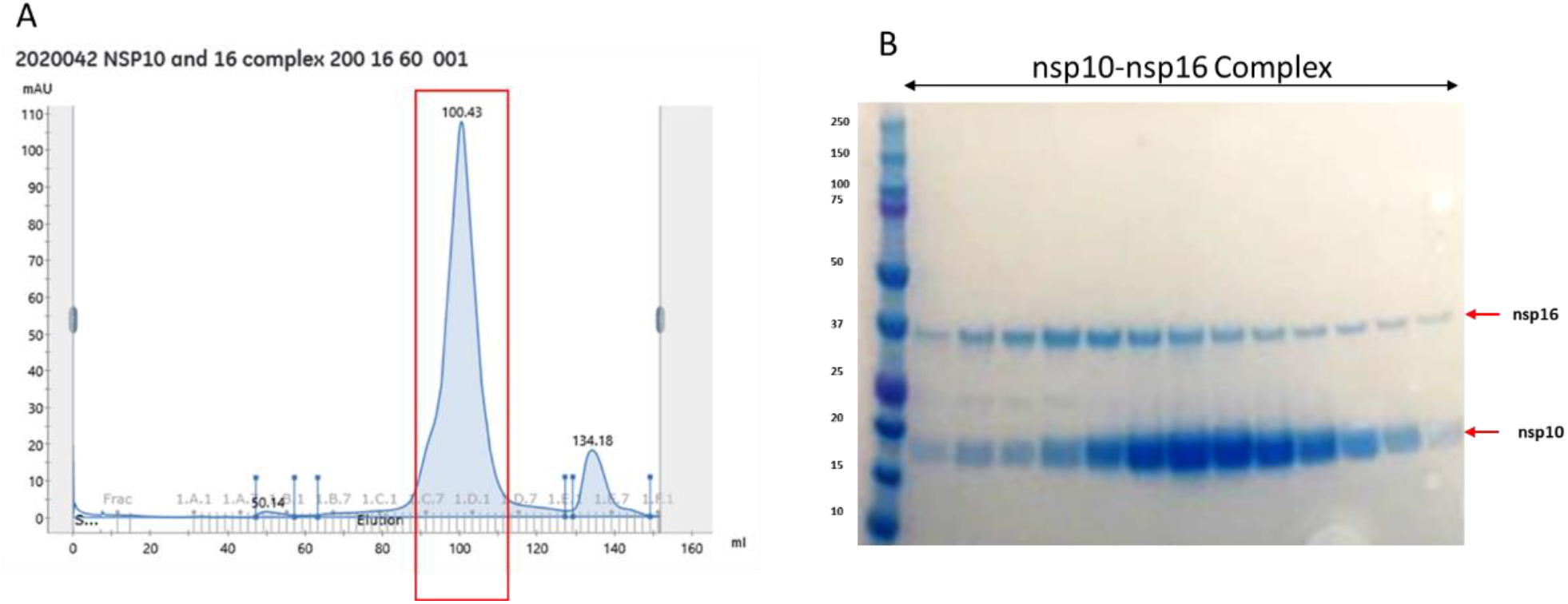
Nsp10-nsp16 complex preparation. SEC was performed on nsp10-nsp16 protein complex using a Superdex 200 16/600 column. (A) nsp10-nsp16 complex SEC profile, (B) SDS-PAGE of the complex fractions resolved by SEC. Protein complex was most stable at 8 (nsp10) to 1 (nsp16) ratio.

**Figure S3.**
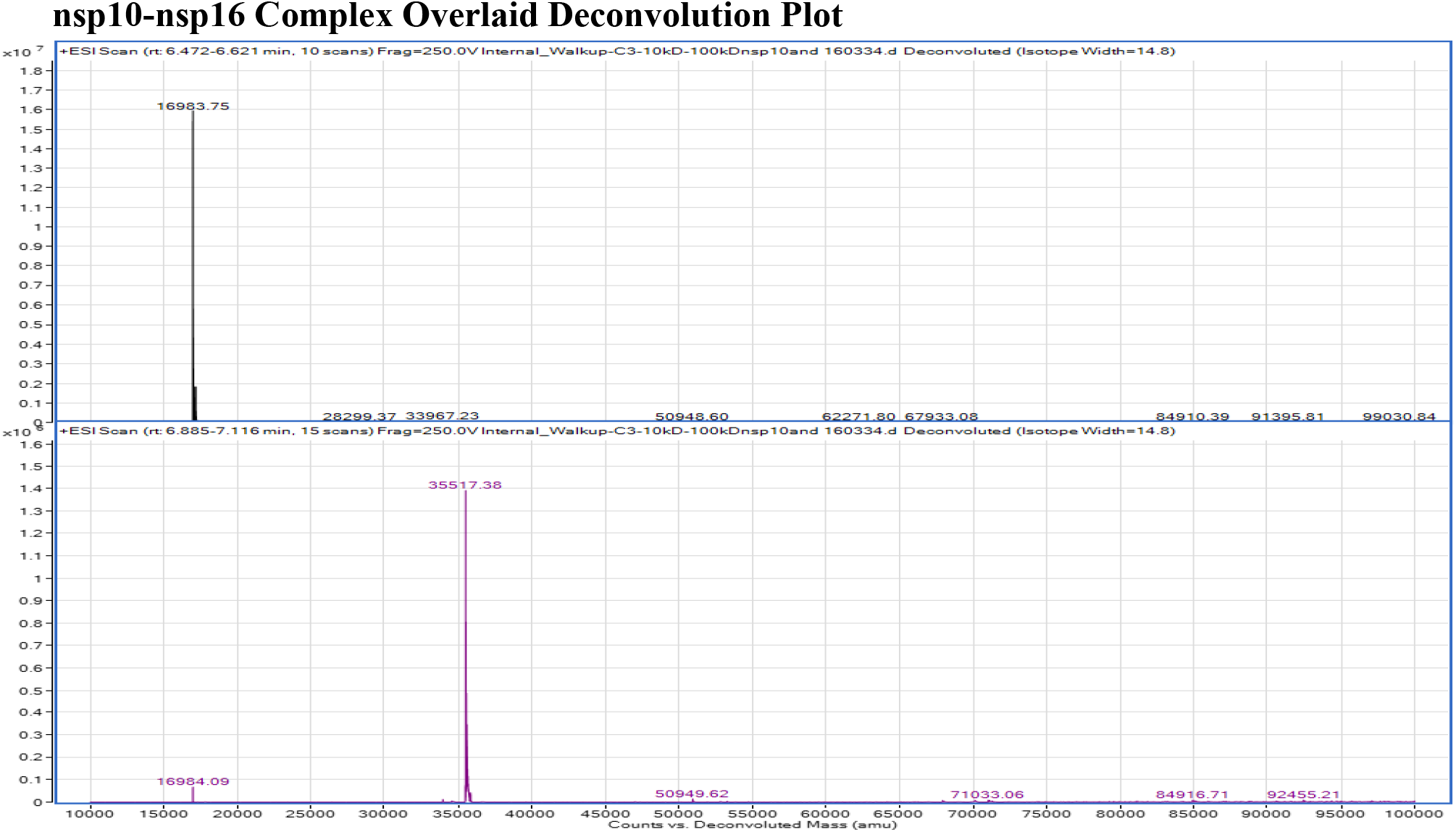
MS profile of the nsp10-nsp16 complex. The MS data confirmed the correct size of the nsp10-nsp16 complex components.

**Figure S4.**
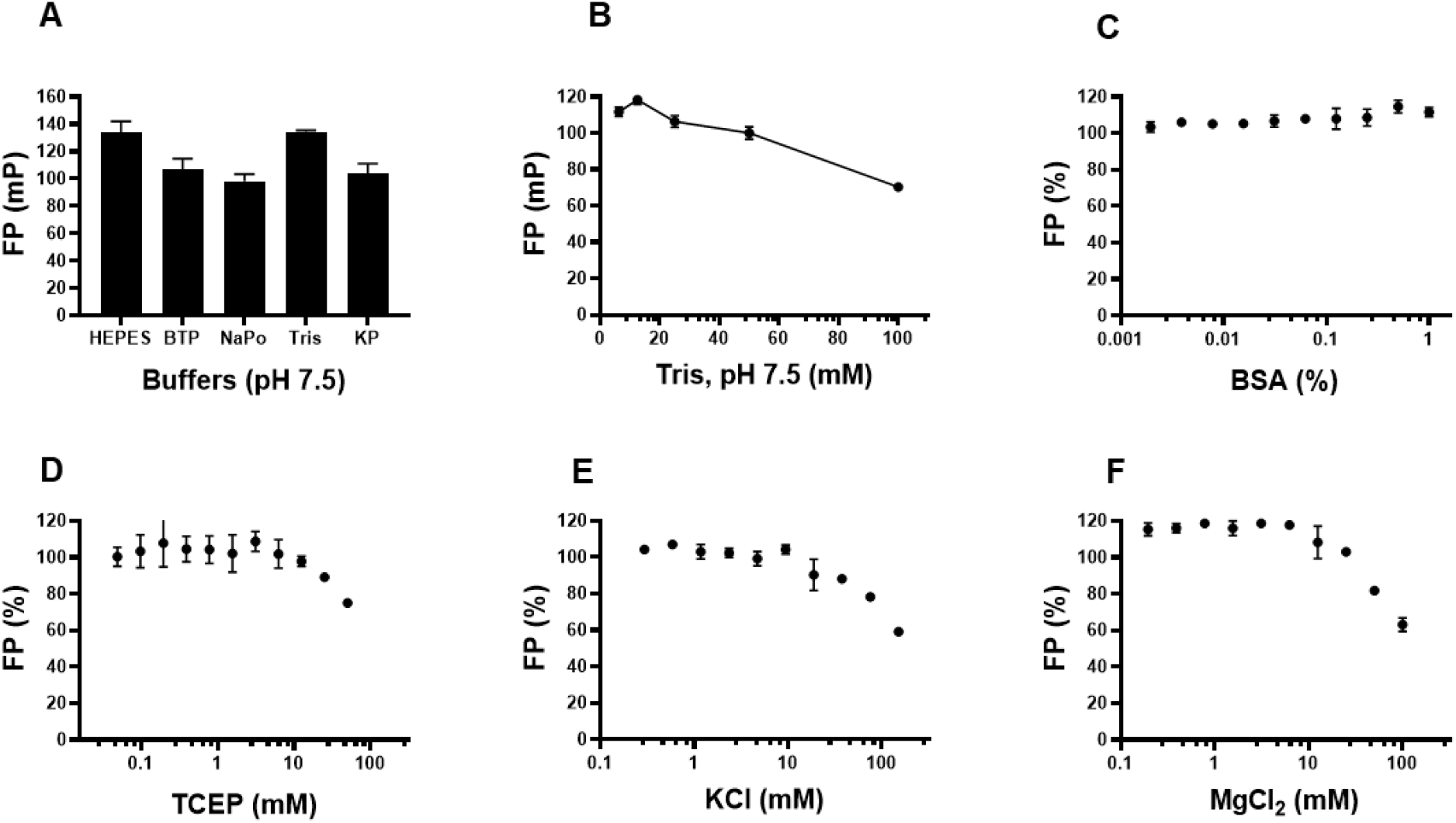
Fluorescence polarization (FP) assay optimization for nsp10-nsp16 complex interaction with N7-meGpppACCCCC-FAM. Fluorescence polarization signal from the interaction of 0.5 μM nsp10-nsp16 with 30 nM of N7-meGpppACCCCC-FAM was evaluated in (A) 20 mM of various buffers; HEPES, Bis-Tris Propane (BTP), Sodium phosphate (NaP), Tris and Potassium phosphate (KP) at pH 7.5, each containing 0.01% Triton X-100; (B) up to 100 mM range of Tris, pH 7.5 and 0.01% Triton X-100; Effect of additives such as (C) BSA; (D) TCEP; (E) KCl; and (F) MgCl_2_ was evaluated in 10 mM Tris, pH 7.5 and 0.01% Triton X-100. Experiments were performed in triplicate (n=3) and plotted values are the average of 3 replicates ± standard deviation. In (A and B), the FP values (mP) are blank subtracted while in (C, D, E and F), FP values are blank subtracted and are presented as the percentage of control (FP %). Data was analyzed using GraphPad Prism software 7.0.4.

**Figure S5.**
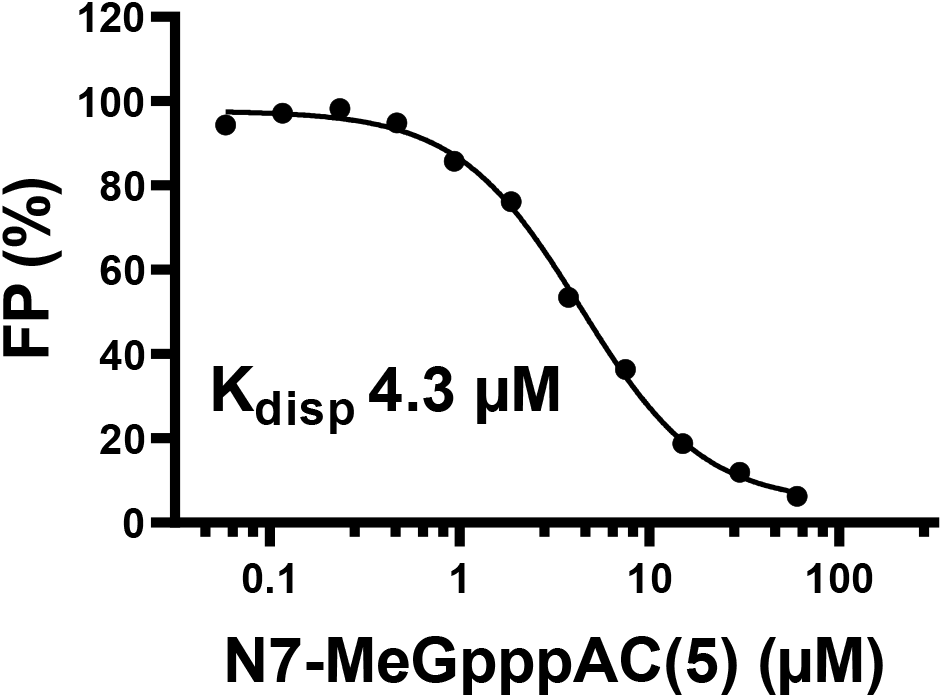
RNA displacement assay monitored by fluorescence polarization. Biotinylated unlabelled RNA (5’ N7-meGpppACCCCC-Biotin 3′; from bioSYNTHESIS) with the same sequence as FAM-labelled RNA (5’ N7-meGpppACCCCC-FAM) displaced the labelled RNA from nsp10-nsp16 complex with a K_disp_ of 4.3 μM and Hill slope of −1.

